# Transcriptional profiling of the cortico-accumbal pathway reveals sex-specific alterations underlying stress susceptibility

**DOI:** 10.1101/2024.10.29.620955

**Authors:** AM Pessoni, L Blanc-Arabe, L Pancotti, S Mansouri, M D’Angelo, K Huot, A Marroquin Rivera, M Peralta, C Zhao, Q Leboulleux, M Lévesque, CD Proulx, B Labonté

**Author notes:** Contributed equally to this work (AM Pessoni, L Blanc-Arabe, L Pancotti, S Mansouri, M D’Angelo).

## Abstract

Anxiety and depressive disorders, including major depressive disorder (MDD), affect millions of people every year, imposing significant socio-economic burdens. In this scenario, current treatments for MDD show limited efficacy, highlighting the need to better understand its molecular mechanisms. The medial prefrontal cortex (mPFC) has been identified as a critical brain region in MDD pathology, displaying altered activity and morphology. This study targets the mPFC-to-nucleus accumbens (NAc) pathway, which is implicated in the regulation of emotional behavior. We used a pathway-specific approach to uncover transcriptional profiles in mPFC neurons projecting to the NAc in stressed male and female mice. Using the RiboTag technique and RNA sequencing, we identified sex-specific gene expression changes, revealing potential roles in stress susceptibility. Differential expression and weighted gene co-expression network analyses revealed distinct transcriptional responses to chronic stress in males and females. Key findings include the identification of the X-linked lymphocyte-regulated 4B (*Xlr4b*) gene, within a highly relevant gene module, as a stress susceptibility driver in males. By experimentally overexpressing the *Xlr4b* gene, we characterized its crucial role in regulating neuronal firing and influencing arborization patterns to promote anxiety-like behavior in a sex-specific fashion. These findings suggest that chronic stress induces unique and shared transcriptional alterations in mPFC neurons projecting to the NAc. Some of these alterations change the morphological and functional properties of neuronal pathways ultimately contributing to the differential manifestation of anxiety-like and depressive-like behaviors in male and female mice.

## Introduction

Anxio-depressive disorders affect 301 million people worldwide^1^. In 2019, major depressive disorders (MDD) became one of the leading causes of disability, with estimates of $1.7 trillion in lost economic productivity^1^. Despite the significant impact MDD has on modern societies, current treatments show only relative efficacy, with roughly one-third of patients showing signs of remission^2,3^. Although more recent therapeutic strategies show encouraging success, the molecular mechanisms underlying the development of the disease remain poorly understood.

Among the brain regions affected in MDD patients, the medial prefrontal cortex (mPFC) has consistently emerged as a hub structure^4–6^. For instance, higher metabolic activity has been reported in the mPFC of MDD patients, and stimulation of this region has been shown to reverse MDD symptoms in treatment-refractory patients while normalizing activity in several brain structures^4,7^. Furthermore, anatomical and morphological analyses reported a lower overall volume and reduced dendritic number and complexity in post-mortem mPFC tissue from human MDD and chronically stressed mice^8–10^. Nevertheless, changes in the activity and morphology of the neuronal populations found in the mPFC alone cannot fully explain the clinical and behavioral manifestations of the disease. These must be understood as changes in the activity of specific neuronal pathways and circuits involving the mPFC.

Recent functional imaging studies confirmed the involvement of the mPFC in networks controlling different behavioral features relevant to the expression of MDD^11–13^. Amongst these structures, the nucleus accumbens (NAc) is known to receive dense projections from the mPFC^14^. Classically, the NAc has been associated with the control of reward, motivation, and goal-directed behaviors^15–17^. More recent functional studies targeting this specific pathway in mice reinforce the idea that it is involved in the control of emotional behaviors^18–21^. For instance, optogenetic stimulation of cortico-striatal neurons modulates reward-seeking behavior^22,23^, and optogenetic stimulation of mPFC glutamatergic terminals in NAc promotes resilience to chronic social defeat stress in male mice^24,25^. Previous studies from our group have also shown that chronic variable stress (CVS) increases the excitability of NAc-projecting mPFC neurons in a sex-specific manner^26,27^. Interestingly, chemogenic activation of this pathway after subchronic stress induces anxiety and behavioral despair in both sexes, while its inhibition rescues this behavioral phenotype in females only^26^. This suggests that increased activity in the cortico-accumbal pathway contributes to the dysregulation of emotional behaviors, although the molecular mechanisms underlying these effects in males and females remain poorly understood.

Transcriptional changes affecting not only gene expression but also the organization of gene networks have been reported across several brain regions and circuits in post-mortem brain tissues of MDD patients and mouse chronic stress models^8,25,27–29^. These changes have been shown to affect the activity of specific neuronal populations in a sex-specific manner^27,30^. For instance, the downregulation of *Dusp6* in the mPFC was shown to increase stress susceptibility by enhancing pyramidal neuron activity in females but not in males^27^. Similarly, the overexpression of *FEDORA*, a long non-coding RNA (lncRNA), in mPFC induces depressive-like behavior by increasing pyramidal neuron excitability in females only^31^. These transcriptional changes, affecting the activity of specific neuronal populations and their respective pathways and circuits, are likely responsible for the expression of distinct behavioral phenotypes. Indeed, precise gene signatures have been associated with the transition from trait to state depression in human mPFC^32^. More recently, our group identified transcriptional signatures across brain regions associated with the expression of specific symptoms of MDD in males and females^29^. Nevertheless, the transcriptional structure of NAc-projecting mPFC neurons, and how it is affected by chronic stress in males and females is still unknown. It is reasonable to speculate that changes in gene expression could explain part of the behavioral phenotype induced by chronic stress by interfering with the intrinsic properties of this neuronal pathway.

In this study, we used a pathway-specific approach to reveal the transcriptional profiles of mPFC pyramidal neurons projecting directly to the NAc in stressed male and female mice. We identified gene signatures and their key master regulators as potential drivers of stress susceptibility in a sex-specific manner by regulating the functional and morphological properties of NAc-projecting mPFC neurons.

## Methods

### Animals

All animal experiments were conducted on male and female mice aged 8 to 14 weeks. The transcriptional experiments were performed on RiboTag mice (B6N.129-Rpl22tm1.1Psam/J; Jackson Laboratory, Bar Harbor, Maine, USA). All other experiments, including *ex vivo* electrophysiology, morphological analyses, and molecular and behavioral validation, were conducted on C57BL/6Ncrl mice (Charles River Laboratories, Saint-Constant, Quebec, CA). All experiments were performed in accordance with the Canadian Guide for the Care and Use of Laboratory Animals and were approved by the Animal Protection Committee of Université Laval. The animals were housed 4 per cage at 22 ± 1°C on a 12-hour light/dark cycle without enrichment. Food and water were provided *ad libitum*.

### Chronic and Subchronic Variable Stress Paradigms

The chronic variable stress (CVS) paradigm was used to induce depressive-like behaviors in both males and females, as previously described^26,27^. Briefly, CVS involves three different stressors repeated over 21 days. On the first day, mice were placed in a shocker and received 100 foot shocks (0.45mA) applied randomly over one hour. On the second day, mice were suspended by the tail for one hour. On the third day, mice were restrained for one hour. Each stressor was repeated daily for a total of 21 days. This procedure consistently induces a complex phenotype defined by anhedonia, behavioral despair, and anxiety^26,27,33^.

A subthreshold version of the chronic variable stress (sCVS) was also used, as described before^34^, to reveal the contribution of target genes to stress susceptibility. By itself, sCVS is insufficient to induce behavioral changes, but can elicit enhanced stress susceptibility when combined with molecular challenges^20,27^. It involves applying the same type of stressors as in CVS but for only 6 days in males and 3 days in females, as 6 days of stress in females is sufficient to drive stress-related phenotypes^20^.

### Behavioral Assessment

A panel of behavioral tests was used to assess anxiety- and behavioral despair-like states. Animals used for transcriptional analyses were not tested to avoid the impact of additional stress induced by behavioral assessment on transcriptional profiles^35^. At the end of sCVS, animals were single-housed, and behavioral tests were performed in the following order: novelty-suppressed feeding test, splash test, and forced swim test. All tests were performed in a dedicated room, separate from the housing and stress rooms. On each test day, mice were habituated to the test room for one hour before testing. Unstressed control groups were handled for 5 minutes for the 6 days prior to the behavioral assessment.

#### Novelty-Suppressed Feeding Test

Novelty-suppressed feeding (NSF) test was used to assess anxiety-like behavior. The test was performed as described previously, with minor modifications^26,36,37^. Animals were food-deprived 24 hours before testing. Mice were placed in the corner of an open field box (50×50×40cm) with corn cob bedding and a single food pellet in the center. Mice were allowed to explore for a maximum of 5 minutes or until they began to eat. They were then immediately transferred to their home cages containing food where all mice ate within 2 minutes. The tests were performed under red-light conditions.

#### Splash Test

The splash test was used to assess depressive-like behavior^26,38,39^. Grooming activity was measured, including nose/face grooming and head and body washing. Mice were sprayed on the back three times with a 10% sucrose solution and placed into an empty transparent acrylic cage under red-light conditions. Their behavior was recorded for 5 minutes, and the total grooming time was hand-scored by an experimenter blind to the experimental conditions.

#### Forced Swim Test

The forced swim test was used to assess the behavioral despair-like state^26,40,41^. Mice were placed in a 3L beaker containing 2L of water at 25 ± 1°C. Their activity was recorded for 6 minutes under white light conditions. Immobility was defined as the absence of struggling, with only the minimal activity necessary to keep the head above water, using a 2-second threshold. The entire video was considered for the assessment of latency to immobility. Animals that didn’t stop swimming after 6 minutes were excluded from the analysis.

#### Emotionality Score

The emotionality score was calculated as previously described^42^. The emotionality score is an average of individual z-scores calculated for each test using the following formula:

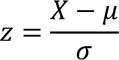

in which *X* represents the individual value for the test, *µ* the control group mean, and *σ* the control group standard deviation. Parameters with inverted correlations, such as grooming time (splash test) and latency to immobility (forced swim test), were converted into positive values to indicate increased emotionality. A global emotionality score was obtained by averaging the z-scores of each behavioral test. Emotionality was calculated separately for male and female animals.

### Viral vectors and stereotaxic injections

All viruses used in this study were generated and obtained from the *Plateforme d’Outils Moléculaires* (POM) at the CERVO Brain Research Centre. AAVs were injected bilaterally into the NAc (coordinates: 10°, anteroposterior: 1.55 mm, mediolateral: ±1.45 mm, dorsoventral: 4.4 mm) and mPFC (coordinates: 10°, anteroposterior: 1.8 mm, mediolateral: ±0.75 mm, dorsoventral: 2.3 mm) of 8-week-old animals. Mice were anesthetized with isoflurane and placed on a stereotaxic frame (Stoelting CO., Wood Dale, IL). Viral vectors were injected at a rate of 0.1 µL/min followed by 5-minute rest. The adeno-associated viruses (AAVs) used in this study are detailed in **Supplemental Table 1.**

For the transcriptional screening experiments, mPFC neurons projecting to the NAc were targeted via the injection of 0.3 µL of AAV2-retro-CAG-Cre-eGFP diluted at 1:2 into the NAc of RiboTag mice. Electrophysiological recordings were performed by injecting 0.3 µL of AAV2-retro-hSyn-Cre diluted at 1:10 into the NAc and 0.4 µL AAV2/5-hSyn-DIO-*Xlr4b*-GSG-T2A-*mCherry* diluted at 1:2 in the mPFC, in wild type mice. Morphological assessments were performed by injecting 0.3 µL of AAV2-retro-CAG-Cre-eGFP diluted at 1:10^3 into the NAc and 0.4 µL of a 1:1 mix of AAV2/5-hSyn-DIO-*Xlr4b*-GSG-T2A-*mCherry* and AAV5-hSyn-DIO-eGFP into the mPFC, in wild type mice. Behavioral tests were performed by injecting 0.3 µL of AAV2-retro-CAG-Cre-eGFP diluted at 1:10 into the NAc and 0.4 µL AAV2/5-hSyn-DIO-*Xlr4b*-GSG-T2A-*mCherry* diluted at 1:2 into the mPFC, in wild type mice. For all experiments, a control group of mice was injected with AAV5-hSyn-DIO-*mChrerry* instead of AAV2/5-hSyn-DIO-*Xlr4b*-GSG-T2A-*mCherry*, using the same volume and dilution.

### RNA sequencing and bioinformatics

#### Tissue collection

Brain tissue from the mPFC was collected from fresh tissue and flash-frozen for storage at −80°C. In total, 21 samples (6 control males, 5 CVS males, 5 control females, and 5 CVS females) were included in this study.

#### RNA extraction, immunoprecipitation, and purification

RNA was extracted, immunoprecipitated, and purified from a portion of each sample, while a smaller portion (5 μL of total volume), used as input controls, was only extracted and purified, as described previously^43^ with minor modifications. Briefly, frozen tissue was thoroughly homogenized in ice-cold CHX-lysis buffer containing 20 mM HEPES–KOH, 5 mM MgCl2, 150 mM KCl, 1 mM DTT, 1% v/v NP40, 200 U/mL RNAse Inhibitor (NEB M0314), 100 μg/mL cycloheximide, and Complete EDTA-free Protease Inhibitor Cocktail. The lysate was centrifuged at 16,000 × g for 10 minutes at 4°C. The input volume was stored, and the remaining supernatant containing the tagged ribosome-mRNA complexes was carefully transferred to a vial containing pre-washed magnetic Protein A/G beads (Thermo Scientific™ Pierce™ Protein AG Magnetic Beads) in CHX-lysis buffer and subjected to gentle head-to-toe rotation in a cold room for 1 hour. The beads were separated from the supernatant using a magnet, and the resulting precleared supernatant was transferred to a new prechilled tube on ice. The precleared lysate was combined with 2.5 μL of polyclonal anti-HA antibody (Abcam, ab9110) and placed in head-to-toe rotation overnight at 4°C. In the following day, washed Dynabeads were added to the mixture and rotated for 4 hours at 4°C. Beads bound to the tagged ribosome-mRNA complexes were isolated from the supernatant using a magnet, washed with buffer containing 20 mM HEPES–KOH, 5 mM MgCl2, 350 mM KCl, 1 mM DTT, 1% v/v NP40, and 100 μg/mL cycloheximide, and finally resuspended in CHX-lysis buffer supplemented with RTL buffer (Qiagen) to allow RNA separation from the beads. The supernatant containing the recovered RNA was collected using a magnet at room temperature. For both fractions, recovered RNA was purified using the RNeasy mini kit (Qiagen) following the manufacturer’s instructions.

#### RNA sequencing

For both fractions, immunoprecipitate (IP) and input, libraries for RNA sequencing were synthesized using Smart-Seq v4 (Takara Bio) at the *Plateforme de séquençage de nouvelle génération* at the *Centre hospitalier universitaire* (CHU) de Québec-Université Laval Research Center. cDNA was made from 85 pg of starting RNA and amplified for 16 PCR cycles. Final dual-index libraries were produced using the Illumina Nextera XT kit following the manufacturer’s recommendations. The quality of final indexed libraries was examined with a DNA screen tape D1000 on a TapeStation 2200 (Agilent Technologies), and quantification was done on a QuBit 3.0 fluorometer (ThermoFisher Scientific). Libraries with unique dual indexes were pooled together and sequenced to a depth of 20 million reads (125 bp paired-end) on a HiSeq 2500 sequencing system (Illumina).

#### Data processing

Sequencing data from IP and input fractions were analyzed using the same criteria, similar to those previously described^44^. Sequencing quality and trim reads were assessed using FASTQC^45^ and Trimmomatic^46^. STAR^47^ was used to align paired-end reads to the GENCODE mouse genome, and R was used for subsequent analyses^48^. All samples included in this study passed the quality assessment. Reads for each sample were counted using featureCounts^49^. A gene was defined as the union of all its exons in any known isoforms, based on GENCODE annotation. Any reads that fell into multiple genes were excluded from the analysis. The threshold for excluding genes expressed at low levels was set to <5 reads in at least 20% of the samples per group, as described previously^27^.

#### Identification of genes not enriched in neurons

As reported previously, immunoprecipitate from RiboTag experiments containing the RNA bound to the tagged complexes may also contain transcripts from genes not expressed in the cell type of interest^44^. To address this, a filter was created based on the expression profile of mouse cell types previously sequenced^50^. Briefly, publicly available mouse RNA-Seq data from purified neurons, astrocytes, microglia, endothelial cells, pericytes, and various maturation states of oligodendrocytes was downloaded (GEO accession ID GSE52564). Next, genes with expression values lower than 1 fragment per kilobase per million mapped fragments (FPKM) were removed, and samples from each cell type were merged using their mean values. A two-step process was implemented to remove genes that were either lowly or not expressed in neurons or enriched in non-neuronal cell types. First, genes with expression values lower than the first quartile (Q1=1.58) in neuronal cells were listed. Second, a ratio between the expression values of neurons and the mean expression values across all other cell types was created. Genes with a ratio lower than the third quartile (Q3=1.55) were considered enriched in other cell types and listed. Finally, the two lists were combined into a final list of 11,032 unique genes that were not enriched in neurons and were removed from the analysis of IP data.

#### Differential expression analysis

Gene expression was normalized using the voom function from the limma package and differentially expressed genes (DEGs) were identified through a generalized linear model (GLM)^51^. As recommended, differential expression analysis was performed first on input vs. IP samples within conditions and then on IP samples across conditions^44^. For the input vs. IP samples analysis, the type of sample (input and IP) and phenotype (stress and control) were included as main factors, and no covariates were included in the analysis design. Genes were considered DEGs (input vs. IP) if their nominal P-value (*p*) was lower than 0.1 in either control or stress. To ensure both statistical and biological relevance, genes that were not differentially expressed in the previous analysis, as well as those from the list of genes not enriched in neurons, were excluded from the IP gene expression table. This resulted in a final expression table of 5173 genes. For the analysis of IP samples across conditions, we included sex (male and female) and phenotype (stress and control) as the main factors, and again no covariates were included in the analysis design. Finally, an individual gene was called differentially expressed if the nominal P-value of its t-statistic was ≤ 0.05.

#### Gene ontology analysis

Gene ontology (GO) analysis was performed using the R package g:Profiler2 on DEGs from male and female mice after CVS, with significant enrichment fixed at FDR < 0.05. The selected terms were obtained from GO and the CORUM database.

#### Transcriptional overlap analysis

To measure the threshold-free transcriptional overlap between male and female mice after CVS, rank-rank hypergeometric overlap (RRHO) analysis was used^52,53^. The gene expression tables were ranked by their P-values and signed based on the fold change. A matrix of hypergeometric P-values was created after evaluating the gene proportion differing from one condition to another. Multiple testing correction was performed using the Benjamini and Yekutieli method^54^, and the obtained values were represented in a heatmap.

#### Gene co-expression network analysis

Weighted gene co-expression network analysis (WGCNA) was employed to identify modules containing genes that exhibit strong co-expression using the WGCNA package (version 1.72.5)^55^. Initially, a sample network was constructed based on Euclidean distance, and a standardized connectivity exclusion threshold was set at −2.5, in which none of the samples were classified as outliers. Subsequently, a unified unsigned network including all 21 IP samples was created, generating a biweight midcorrelation matrix between all gene pairs, which was transformed through a few steps: first, it was converted into an unsigned adjacency matrix using a specific soft threshold power and then transformed into a topological overlap matrix^56,57^. Using average linkage hierarchical clustering, groups of genes were created, and a subsequent dynamic tree cut was employed to investigate clusters in a nested dendrogram, identifying modules of highly co-expressed genes. Each module was assigned a unique arbitrary color label for characterization and associated with a significant gene ontology term (containing at least three related genes, when possible). To highlight genes with significant intramodular connectivity, the top 5% of genes within each module were designated as hub genes.

#### DEGs enrichment, phenotype association, and relevance score

Module differential expression enrichment for genes significantly upregulated, downregulated, or both was assessed in each module for both sexes, using the GeneOverlap package from Bioconductor, considering 5,173 genes as the whole set. Fisher’s exact tests (FET) were used to perform all enrichment assessments, with significance fixed at nominal P-value ≤ 0.05. Phenotype association with network modules was performed by calculating t-tests for the module eigengene (ME) values between stress and control samples of each sex, with significance considered at nominal P-value ≤ 0.05. Gene significance (GS), the gene association with phenotype, was calculated with point-biserial correlations between stress and control samples of each sex, only for genes with average expression values greater than 1, with significance considered at nominal P-value ≤ 0.05. The module relevance score was calculated by combining individual scores attributed to different ranges of P-values obtained in the analysis. DEG enrichment scores were assigned for three categories (upregulated, downregulated, and both) based on the following ranges: *p* ≥ 0.05 = 0; 0.05 > *p* ≥ 0.001 = 1; 0.001 > *p* ≥ 0.0001 = 2; 0.0001 > *p* ≥ 0.00001 = 3; and *p* < 0.00001 = 4. Additionally, these scores were adjusted based on the odds ratio (OR), with values added as follows: OR ≤ 1 = 0; 1 < OR ≤ 2 = 1; 2 < OR ≤ 4 = 2; 4 < OR ≤ 10 = 3; 10 < OR = 4. Phenotype association scores were attributed as follows: *p* ≥ 0.05 = 0; 0.05 > *p* ≥ 0.01 = 1; 0.01 > *p* ≥ 0.001 = 2; 0.001 > *p* ≥ 0.0001 = 3; and *p* < 0.0001 = 4. Both males and females were given individual scores, and the relevance of each module was ranked based on the scores from the male samples.

#### Cytoscape visual representation

The network structure of selected modules was visualized using Cytoscape (version 3.10.1)^58^. Briefly, node and edge values were obtained for the connections and exported as input for Cytoscape. Style parameters were then adjusted to highlight target genes, hub genes and their connections, and genes differently expressed in males or females.

### Electrophysiological recordings

Whole-cell patch-clamp experiments were performed as previously described^59^. Briefly, sCVS was performed three weeks after viral injections in male mice. After 6 days of sCVS, mice were anesthetized with isoflurane and received transcardiac perfusion with 10 mL of ice-cold NMDG-artificial cerebrospinal fluid (aCSF) solution containing (in mM): 1.25 NaH2PO4, 2.5 KCl, 10 MgCl2, 20 HEPES, 0.5 CaCl2, 24 NaHCO3, 8 D-glucose, 5 L-ascorbate, 3 Na-pyruvate, 2 thiourea, and 93 NMDG (osmolarity adjusted to 300–310 mOsmol/L with sucrose). The pH was adjusted to 7.4 using 10 N HCl. Kynurenic acid (2 mM) was added to the perfusion solution on the day of the experiment. The brains were then quickly removed, and 250 μm acute brain slices containing the mPFC were prepared using a Leica VT1200S vibratome. The slices were placed in a 32°C oxygenated perfusion solution for 10 min and were then incubated for 1 h at room temperature in HEPES-aCSF solution (in mM): 1.25 NaH2PO4, 2.5 KCl, 10 MgCl2, 20 HEPES, 0.5 CaCl2, 24 NaHCO3, 2.5 D-glucose, 5 L-ascorbate, 1 Na-pyruvate, 2 thiourea, 92 NaCl, and 20 sucrose (osmolarity adjusted to 300–310 mOsmol/L with sucrose). The pH was adjusted to 7.4 using 10 N HCl. The slices were then transferred to a recording chamber on the stage of an upright microscope (Zeiss), where they were perfused with 3-4 mL/min of aCSF (in mM): 120 NaCl, 5 HEPES, 2.5 KCl, 1.2 NaH2P04, 2 MgCl2, 2 CaCl2, 2.5 glucose, 24 NaHCO3, and 7.5 sucrose. The perfusion chamber and the aCSF were maintained at 32°C. All solutions were oxygenated with 95% O2/5% CO2. A 60x water immersion objective and a video camera (Zeiss) were used to visualize neurons in the mPFC. Fluorescent neurons were visualized by transmitting 568 nm light through the microscope light path with a LED colibri set (Zeiss). Recording pipettes made of borosilicate glass (3-7 MΩ resistance) were pulled using a P-1000 Flaming/Brown micropipette puller (Sutter Instruments). Recordings were performed using a Multiclamp 700B amplifier (Molecular Devices). For the current-clamp recordings, the intracellular solution consisted of (in mM): 130 K-gluconate, 5 KCl, 10 HEPES, 2.5 MgCl2, 4 Na2ATP, 0.4 Na3GTP, 10 Na-phosphocreatine, and 0.6 EGTA (pH 7.35). Signals were filtered at 5 kHz using a Digidata 1500B data acquisition interface (Molecular Devices, San Jose, CA) and acquired using pClamp 10.6 software (Molecular Devices). For intrinsic properties, neurons were subjected to 12 current steps, from −80 to +140 pA (Δ20 pA). For the Intensity-Frequency (I-F) graph, positive current steps were applied, up to 400 pA (Δ50 pA). After recordings were completed, the slices were fixed in 4% formaldehyde for 24 hours and then transferred to a 0.1M phosphate buffer solution for *post hoc* histological analysis. Electrophysiological traces were analyzed with Clampfit (pClamp Suite, Molecular Devices) and Easy Electrophysiology V2.5.

### Morphological analyses

sCVS was performed three weeks after viral injections in male mice. After 6 days of sCVS, both stress and control mice were anesthetized with a 20% urethane solution injected intraperitoneally. The mice were then perfused with an ice-cold phosphate-buffered saline (PBS 1X) followed by a filtered ice-cold paraformaldehyde (PFA 4%) to wash the residual blood and fix the tissue. The brains were collected and post-fixed in PFA 4% overnight. Brains were cut into 100µm-thick slices using a vibratome (Leica VT1000S, Leica Biosystems) and placed in PBS 1X. The mPFC slices were mounted on glass slides with a protective medium (ProLong Gold Antifade Reagent, Invitrogen, #P36930).

Image acquisition was performed with a confocal microscope (Nikon A1R HD) using a 60X oil immersion objective. Stacked images of the pyramidal neurons labeled with both fluorophores, *mCherry* and eGFP, in the mPFC region were captured with 1µm between each stack. Morphological 3D reconstruction of the neurons was performed using neuTube^60^, and Sholl analysis was performed with Fiji^61^ starting at 15 µm of the soma with 5µm intervals.

For spine density analyses, the image acquisition for the 100 µm thick slices was performed as previously described, using the same stacked images utilized for morphological analysis, merged into a single plane. Each reconstructed neuron was overlaid with those from the merged image to correctly identify the projections and allow for spine density calculation. To determine spine density, we used the Dendritic Spine Counter Fiji plug-in^61^. After manually defining the edges of a spine, every branch belonging to a single neuron was tracked. The plug-in then automatically detected and counted spines. Finally, the operator performs a double verification, with local missing spines manually added to the original count.

### Statistical analyses

Although sample size calculations were not performed, the sample size in this study is justified based on several previously published reports using similar or even smaller sample sizes and showing the power to detect significant statistical differences. A preestablished alpha value of 0.05 was considered for hypothesis testing. Details of each analysis are provided in the respective section.

#### Transcriptional analyses

Samples from 21 animals were used for the transcriptional analysis, including 6 control male, 5 CVS male, 5 control female, and 5 CVS female samples. No outliers were removed from the differential expression analysis or weighted co-expression analyses. After all filters, the final gene expression table used for all transcriptional analyses included 5173 genes enriched in neuronal cells. Differential expression was not corrected for multiple testing but network analysis, including network construction and GO enrichment, was corrected for multiple testing. Details of each analysis are provided above in the respective sections.

#### Behavioral test analysis

64 animals, divided into two cohorts, were included in the behavioral analysis (32 males: 9 *mCherry* control, 7 *mCherry* CVS, 10 *Xlr4b* control, 6 *Xlr4b* CVS; 32 females: 8 *mCherry* control, 7 *mCherry* CVS, 8 *Xlr4b* control, 9 *Xlr4b* CVS). 12 animals were removed due to imprecise viral infections. Outliers were identified using the ROUT method (Q=1%) and 2 outliers were removed (1 male *mCherry* CVS, 1 female *mCherry* CVS). To avoid cohort effects, raw data were standardized using the following formula:

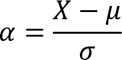

where *α* represents the final standardized value, *X* represents the individual value for the test, *µ* the cohort mean for the test, and *σ* the cohort standard deviation for the test. Two-way ANOVA analysis was performed with *‘viral approach’* and *‘stress’ as main factors.* Tukey *post hoc* analysis was also conducted for multiple comparisons. Statistical analyses were performed using GraphPad Prism 10 (GraphPad, San Diego, CA).

#### Electrophysiology

In total, patch-clamp recordings were obtained from 91 cells of 16 animals (*mCherry* CTRL: 4 animals, 19 cells; *mCherry* sCVS: 4 animals, 21 cells; *Xlr4b* CTRL: 4 animals, 23 cells; *Xlr4b* sCVS: 4 animals, 28 cells). For the I-F analysis, a linear mixed-effects model was performed, with *‘applied current’*, *‘viral approach’* (representing the effect of gene, *mCherry*/*Xlr4b*), and *‘stress’* (CTRL/sCVS) as the fixed effects. The interactions between pairs of variables and a random effect for each cell were included in the model. Tukey *post hoc* analysis was conducted for multiple comparisons. For the other analyses, a Two-Way ANOVA analysis was performed, with *‘viral approach’* and *‘stress’* as the main factors. Tukey *post hoc* analysis was also conducted for multiple comparisons. Statistical analyses were performed using Prism 10 (GraphPad Software) and the R environment.

#### Morphology

Given that the measurement scale of the number of intersections (complexity) corresponds to a discrete count and that it is obtained in a way the measurements correspond to repeated measures clustered within a unique cell, it was deemed appropriate to implement a Poisson generalized linear mixed model (GLMM). First, the distribution of the dependent variable was assessed to verify the appropriateness of using a Poisson GLMM, according to graphical examination (quantile-quantile plots). Subsequently, an intercept-only model was evaluated, gradually adding fixed and random effects. The most parsimonious model was selected by comparing full models (with interactions, random slopes, and random intercepts) with reduced nested models through the Likelihood ratio test (LRT). When differences between the deviance values were not significant, the reduced model was retained; otherwise, the full model was kept. Predictors were centered, and marginal and conditional coefficients of determination were estimated following the Nakagawa and Schielzeth method^62,63^. Statistical analyses were conducted using the R environment, specifically the lmer4, sjPlot, ggplot2, MASS, and lmerTest packages^48,64,65^. For the spine density analysis, a Two-Way ANOVA was performed, with ‘*stress*’ (CTRL/sCVS) and ‘*viral approach*’ (*Xlr4b*/*mCherry*) as the main factors. Tukey *post hoc* analysis was conducted for multiple comparisons.

## Results

The main objective of this study was to evaluate the extent of the stress-induced reorganization of transcriptional profiles in mPFC neurons projecting to the NAc and determine the behavioral, functional, and morphological consequences of these changes in male and female mice. Using a trans-sectional viral approach in RiboTag mice, we mapped transcriptional signatures from cortico-accumbal neurons, performed weighted gene co-expression network analyses, and identified key candidate molecular drivers of stress susceptibility in males and females. We next used viral-mediated gene transfer in mPFC neurons projecting to the NAc to evaluate the impact of this transcriptional reorganization at the behavioral level. Finally, we used electrophysiological and morphological approaches to evaluate the effects of target gene overexpression on the functional and morphological properties of neurons in the cortico-accumbal pathway. Our findings suggest that chronic stress induces unique and shared transcriptional alterations in NAc-projecting mPFC neurons, with important implications for understanding the molecular basis of stress-related behaviors and neuronal function.

### Global Reorganization of Transcriptional Profiles in Cortico-Accumbal Neurons of Stressed Males and Females

In order to define the sex-specific transcriptional and molecular pathways through which CVS impacts NAc-projecting mPFC neurons, we used a RiboTag approach to isolate RNA from mPFC neurons projecting directly to the NAc. This technique leverages the unique feature of the RiboTag mouse, which carries a ribosomal protein gene (Rpl22) with a floxed C-terminal exon followed by an identical exon tagged with hemagglutinin (HA)^43^. We infected RiboTag mice with a retrograde AAV expressing cre recombinase (AAV2-retro-hSyn-Cre-eGFP) in the NAc to allow the expression of the HA-Rpl22 in mPFC neurons projecting specifically to the NAc (**Figure 1A**). After stressing the animals for 21 days according to our CVS protocol, we confirmed the co-localization of HA-Rpl22 and cre-GFP in NAc-projecting mPFC neurons (**Figure 1B**). We then immunoprecipitated the polyribosomes tagged with HA and synthetized RNAseq libraries for sequencing following established protocols^44,66^.

**Figure 1.**
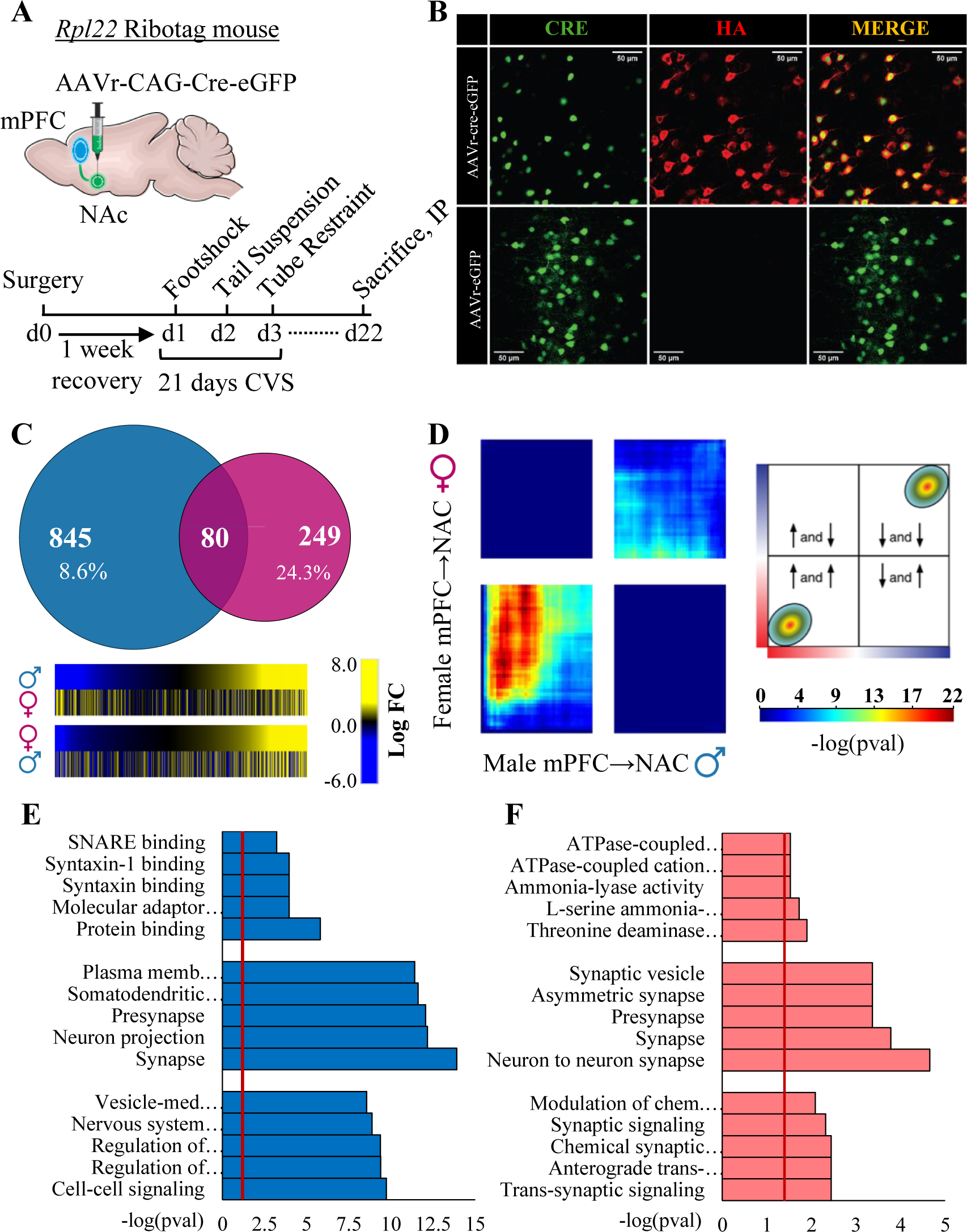
Differential gene expression patterns diverge in male and female mice after CVS. **A.** Schematic representation of a mouse brain showing the mPFC and NAc pathway connections, the viral injection sites, and the experimental timeline for CVS. **B.** Representative example of cre-dependent HA expression (red) due to the injection of AAVr-hSyn-Cre-eGFP (green). **C.** Venn diagram of DEGs (*p* ≤ 0.05) showing low transcriptional overlap between male (blue) and female (pink) mice after CVS, and heatmaps comparing transcriptional changes in male and female mice after CVS. The first pair of heatmaps shows the direction of changes in females for genes that are significantly differentially expressed in males, while the second pair of heatmaps shows the direction of changes in males for genes that are significantly differentially expressed in females. **D.** RRHO heatmaps showing threshold-free transcriptional overlap between male and female mice after CVS. The lower left and upper right quadrants represent an overlap for concomitantly upregulated and downregulated genes, respectively, and the other two quadrants represent the overlap of genes with opposite directions in expression. The color bar represents the degree of significant overlap between transcriptional signatures. **E, F.** Top 5 significantly enriched (*p*_adj_<0.05) gene ontology terms of each category, molecular function, cellular component, and biological process, respectively from top to bottom, related to the DEGs of male (blue) and female (light red) mice after CVS.

We first conducted a differential expression analysis to identify genes significantly up or downregulated (*p*≤0.05) in mPFC neurons projecting to the NAc in stressed male and female mice. Our analysis revealed numerous genes differentially expressed in both males and females, with minimal overlap between both sexes (**Figure 1C, Supp. Table 2**). Specifically, only 8.6% (n=80) of the differentially expressed genes in stressed males were also differentially expressed in stressed females, which was consistent with the different directionality observed in stressed males and females. Together, these finding align with previous reports on bulk tissue in stressed mice and humans with MDD^27,29,30,67,68^ and suggest that stress impacts the transcriptional programs of NAc-projecting mPFC neurons differently in males and females.

We next used a rank-rank hypergeometric overlap analysis (RRHO) to compare male and female transcriptional signatures without imposing stringent statistical thresholds^52,53^. This technique ranks the gene lists by differential expression P-values and takes into consideration the fold change direction, displaying the significance of the overlap intensity^52^. Interestingly, our results revealed a notable overlap between stressed male and female mice (maximum −log_(P-value)_=22), mainly for genes commonly upregulated, but also for downregulated genes in a smaller magnitude (**Figure 1D**). Similarly, a comparison of the top gene ontology terms enriched with DEGs in males and females identified similar biological processes in both sexes related to synapse signaling and trans-synaptic transport, as well as shared cellular components such as synapse and vesicle-related terms (**Figure 1E, F**). In particular, the top terms found in female DEGs were highly focused on synaptic activity, some of which were common in stressed males. Male DEGs exhibited a broader range of terms related to brain activity, with fewer terms shared in females (**Supp. Figure 1A**). Overall, this suggests that while transcriptional signatures in males and females may be similarly affected by stress, there is low overlap when using stringent statistical thresholds. This implies that distinct transcriptional changes in males and females may lead to similar functional alterations in both sexes.

Finally, we evaluated the specificity of our findings in cortico-accumbal neurons by comparing our pathway-specific data with those obtained from bulk sequencing of the whole mPFC in males and females after CVS^27^. Interestingly, our data detected only a minimal number of DEGs commonly affected in both datasets (**Supp. Figure 1B**). This was true for both males and females, with a very small overlap between the sexes. However, a larger overlap was observed with RRHO (although modest: maximum −log_(P-value)_=5), revealing genes commonly up and downregulated in both our bulk and pathway-specific datasets, mostly in males (**Supp. Figure 1C**). Together, these findings suggest that CVS induces a significant proportion of unique transcriptional alterations in mPFC neurons projecting to the NAc, alongside broader changes affecting mPFC neurons more generally in males and females.

### Stress changes the transcriptional organization of the cortico-accumbal transcriptional profile in males and females after CVS

We next used weighted gene-co-expression analysis (WGCNA) to create a unique gene network specific to mPFC neurons from the cortico-accumbal pathway in males and females (**Supp. Figure 2A**). This strategy uses a correlation network to describe pairwise relationships between genes, allowing the creation of gene clusters (modules) composed of correlated nodes, and identification of highly connected nodes (hubs) that can represent that module and significant modules related to a feature of interest^55^. With the aim to evaluate the effect of chronic stress on the transcriptional organization of gene networks in this neuronal pathway, this analysis allowed us to compare directly males and females, evaluating the impact of stress in a sex-specific fashion. In total, we identified 36 modules enriched with genes relevant to specific ontological terms related to neuronal functional domains (**Supp. Figure 2A, Supp. Table 3**). We then calculated the module *eigengene* (ME), which is the global representation of gene variation inside a module, for each sample by extracting the first principal component value from each identified module. This allowed us to calculate the association of specific modules with the expression of a stress phenotype in males and females, distinctly, as previously performed^29,55^. In total, we identified 8 modules associated with stress (*p*<0.05) in males and 9 in females and 3 modules with a trending association (0.05<*p*<0.1) in males and 6 in females (**Figure 2A**). Overall, this represents 30.5% of the modules in males and 41.7% in females, respectively, showing signs of association with the phenotypes and potentially underlying some of the functional and behavioral impact of stress in both sexes.

**Figure 2.**
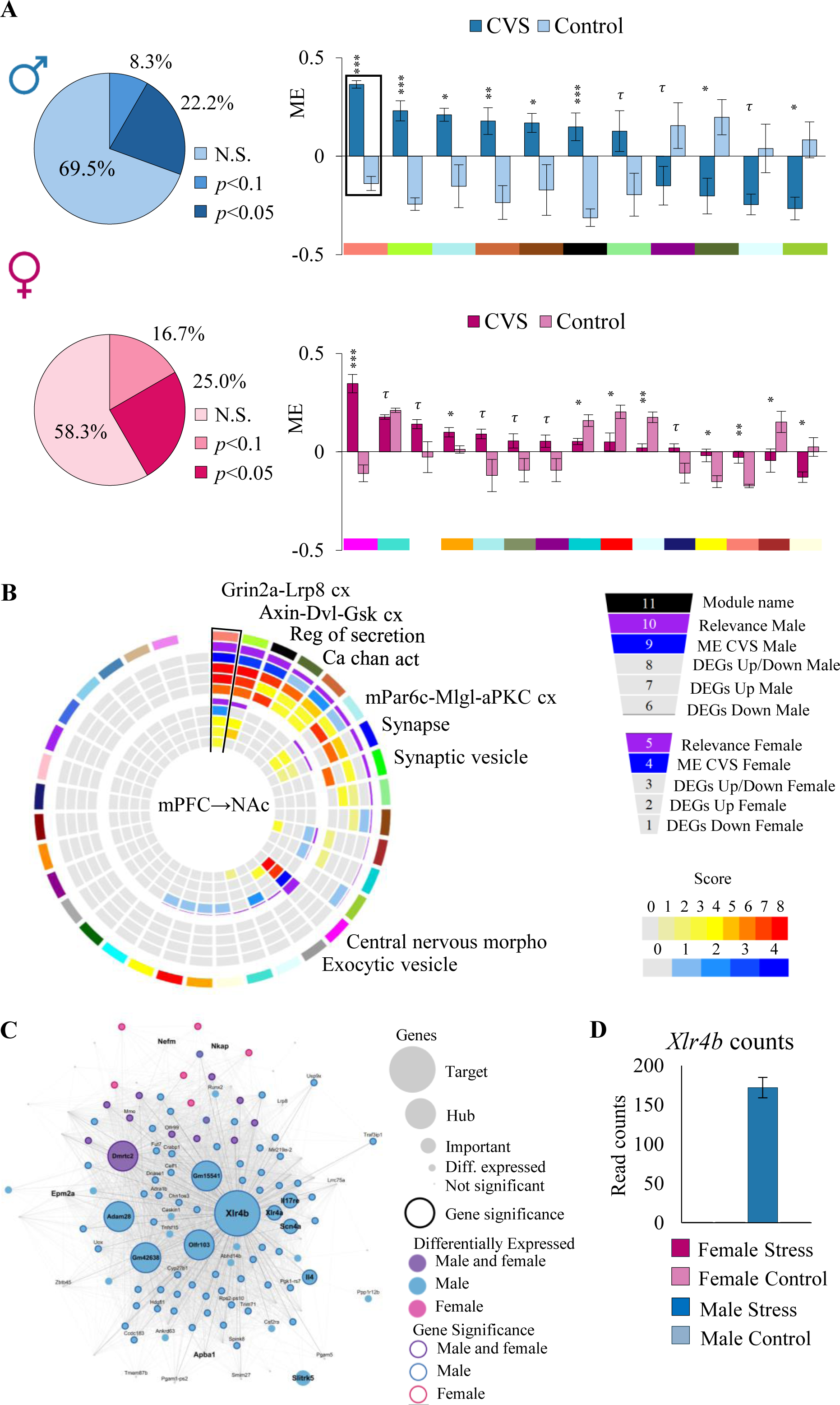
Gene coexpression network from male and female mice after CVS shows modules enriched for DEGs and significantly associated with the phenotype. **A.** Proportion of network modules significantly associated with the stressed phenotype in males (blue) and females (pink), categorized by *p*: ≤ 0.1, ≤ 0.05, and non-significant enrichment, and comparison of ME values between CVS and control mice for males (blue) and females (pink), focusing on the significant (*p* ≤ 0.05) and trending (0.1 > *p* > 0.05) network modules. **B.** Circos plot displaying gene coexpression modules ranked by their relevance score in males (purple). The relevance is measured by the association of ME values with the phenotype (blue) and the degree of enrichment for DEGs (red), whether they are upregulated, downregulated, or both, considering their odds ratios. The outer layers represent male modules, and the inner layers represent female modules. Each module is uniquely characterized by an arbitrary color, shown in the outermost rectangle. Some modules are also labeled with their most significant gene ontology term (*p*_adj_<0.05). **C.** Cytoscape representation of the *Salmon* gene module, the most relevant module in males and second most relevant in females, representing genes differently based on their connectivity, expression, and significance. **D.** Bar plot showing the sum of raw counts for all samples of each sex and condition, highlighting the difference between the expression of the *Xlr4b* gene in male stressed samples when compared to every other.

We tested whether male and female gene modules were enriched for DEGs following chronic stress. In males, 12 modules were enriched with DEGs, either up and/or downregulated, including some related to *Grin2a-Lrp8 complex* (*Salmon*: *p*_adj_<0.01), *Calcium channel activity* (*Dark olivegreen*: *p*_adj_<0.05) and *Function of the synapse* (*Blue*: *p*_adj_<0.001; **Figure 2B, Supp. Table 3**). In females, 8 gene modules were found enriched with either up and/or downregulated genes including the *Turquoise* module, associated with *Behavior* (*p*_adj_<0.01), *Magenta*, enriched with genes related to *Morphology of the central nervous system* (*p*_adj_<0.05), and *Grey60*, relevant to *Exocytic vesicle* (*p*_adj_<0.05; **Figure 2B**, **Supp. Table 3**). Finally, module association with the phenotype and DEG enrichment values were used to compute a relevance score (see material and methods) which allowed to rank modules the most importantly associated with the effect of chronic stress in males and females (**Figure 2B**).

Among the top candidate gene modules, the module *Salmon* ranked as the most relevant in stressed males (*relevance score*=26) and the second most relevant in stressed females (*relevance score*=12; **Figure 2A**, **B**, **Supp. Table 3**). This module comprises 136 genes and is significantly enriched for genes associated with the *Grin2a-Lrp8 complex* (*p*_adj_=0.0096). It is enriched for genes up (*p*=5.54×10^−33^ for males and *p*=0.026 for females) and downregulated (*p*=6.32×10^−6^ for males and *p*=0.008 for females) and for genes associated with stress phenotype (*p*=2.91×10^−45^) in males but not in females (p=0.13; **Supp. Table 3**). It is also associated with the expression of stress responses in both sexes (*p*=2.94×10^−6^ for males and *p*=0.0068 for females; **Figure 2B**, **Supp. Table 3**). In fact, 68.38% (N=93) of genes composing this module (*Salmon)* are associated with the expression of stress responses in males and 13.97% (N=19) in females. The *Salmon* module is composed of 6 hub genes, including *Xlr4b*, *Adam28*, *Gm15541*, *Gm42638*, *Dmrtc2*, and *Olfr103* (**Figure 2C**). Importantly, all these hub genes are differentially expressed in stressed males, while only *Dmrtc2* was also downregulated in stressed females. Similarly, each hub is significantly associated with the phenotype in males, while only *Dmrtc2* associates with the phenotype in females (**Figure 2C, Supp. Table 2**). Hubs are genes highly correlated with the module eigengene^55,69^ and, consequently, with the expression of a larger number of genes within their respective modules. Thus, hub genes are believed to play important roles in the structure of gene networks with downstream functional and behavioral consequences^25,27,70,71^. As illustrated in **Figure 2C**, the graphical representation of this network highlights key genes within the module, including hub and node genes. Our analysis revealed that the X-linked lymphocyte-regulated 4B (*Xlr4b*) gene is a primary contributor to the module’s association with stress expression in male mice. *Xlr4b* is amongst the most differentially expressed genes (*p*<5.0×10^−9^) with the highest fold change (FC=8.28) in stressed males versus controls, with no change in stressed females (**Supp. Table 2**). It is also significantly associated with the expression of the phenotype in males (*p*=1.98×10^−12^) and its expression is strongly induced by stress in males, while undetected in control and female mice (**Figure 2D**).

Besides, we also identified gene modules associated with unique characteristics and potential implications for stress responses in either males or females. In females, we identified The *Magenta* module as the gene module most significantly associated with the effect of stress (*p*=9.47×10^−5^). This module is composed of 163 genes relevant to *Central nervous system morphogenesis* (*p*_adj_=0.0311; **Figure 2B**). Importantly, the *Magenta* module is enriched for upregulated genes (*p*=7.56×10^−36^) and with genes associated with stress phenotype (*p*=1.23×10^−13^) in females but not males (*p*=0.99; **Supp. Table 3**). It contains 8 hub genes including *Vamp9*, *Gm37311*, *4933423P22Rik*, *Gm47572*, *Gm6117*, *Zfp108*, *E430024I08Rik*, and *Zfp109*, which are also differentially expressed in stressed females exclusively (**Supp. Figure 2B**).

In males, our analysis identified the *Black* module as significantly associated with the effect of stress (*p*=8.68×10^−4^; **Sup. Table 3**). This module comprises 197 genes enriched in functional terms related to *cell-cell signaling* (*p*_adj_=0.0118; **Figure 2B**). It is significantly enriched for up (*p*=4.51×10^−4^) and downregulated (*p*=1.58×10^−17^) genes in males but not females (**Sup. Table 3**). It is also enriched for genes significantly associated with the phenotype in males (*p*=9.84×10^−25^) but not females (*p*=0.89). The *Black* module includes 9 hub genes including *Gata1*, *Gm15179*, *Gm23368*, *Gm7478*, *Gm26802*, *Utf1*, *Pgrmc1*, *Tmem59l*, and *Smt3h2-ps4*, which are all significantly associated with the phenotype and differentially expressed in stressed males specifically (**Supp. Figure 2C**). Additional modules relevant to the effects of stress in both males and females are provided in **Supp. Table 3**. Overall, this suggests that chronic stress changes the transcriptional organization of gene networks in mPFC neurons in the cortico-accumbal pathway in a sex-specific fashion. Interestingly, while our analysis identified modules relevant for virtually every aspect of neuronal activity and functional properties, only a proportion of them are affected in males and females.

### *Xlr4b* overexpression in NAc-projecting neurons coupled with sCVS induces anxiety- and despair-like behavior only in male mice

Our analyses highlighted the *Salmon* gene module and its hub gene *Xlr4b* in mPFC neurons projecting to the NAc as main drivers of stress susceptibility in males. Interestingly, *Xlr4b* was shown to control dendrite and spine formation through a Cux1/2-dependent mechanism^72^. Variations in *Xlr4b* expression have also been associated with working memory and affective deficits in mice^72,73^ and autism and schizophrenia in humans^74–76^. Consequently, we hypothesized that modulating the activity of the *Salmon* module by reproducing the stress-induced expression pattern of *Xlr4b* in mPFC neurons from the cortico-accumbal pathway would change the male’s susceptibility to stress.

We started by testing whether the overexpression of *Xlr4b* alone would be sufficient to induce anxiety and depressive-like behaviors. *Xlr4b* was specifically overexpressed in mPFC neurons projecting to the NAc using a trans-sectional viral approach in both males and females (**Figure 3A, B, G, H)**. Our analyses suggest that *Xlr4b* overexpression in NAc-projecting mPFC neurons alone does not trigger any behavioral modification in male and female mice (**Figure 3C-F, I-L**). Importantly, the expression of complex stress phenotypes can be triggered when stress-relevant molecular features are reproduced in conjunction with the administration of subthreshold stress procedures which, by themselves, do not induce any behavioral alterations^20,27^. Accordingly, we next tested whether the overexpression of *Xlr4b*, combined with subthreshold stress (sCVS), triggers the expression of anxiety and depressive-like behaviors in males and females. Our results show that *Xlr4b* overexpression in cortico-accumbal mPFC neurons, combined with 6 days of CVS in males and 3 days in females, significantly increased the time before eating in the NSF test (F_(1,28)_=13,242, *p*=0,001) and reduced the grooming time in the splash test (F_(1,28)_=8,597, *p*=0,007) in male (**Figure 3C, D**) but not female mice (**Figure 3I, J**), with no effect observed in the forced swim test in both males and females (**Figure 3E, K**). Interestingly, *Xlr4b* overexpression combined with sCVS also significantly increased overall behavioral emotionality in males (F_(1,28)_=6,414, *p*=0,017; **Figure 3F**) and, surprisingly, slightly reduced it in female mice (F_(1,29)_=1,214, *p*=0,019; **Figure 3L**). Overall, these results suggest that the overexpression of *Xlr4b* in mPFC neurons from the cortico-accumbal pathway drives stress and emotional susceptibility in males but not females.

**Figure 3.**
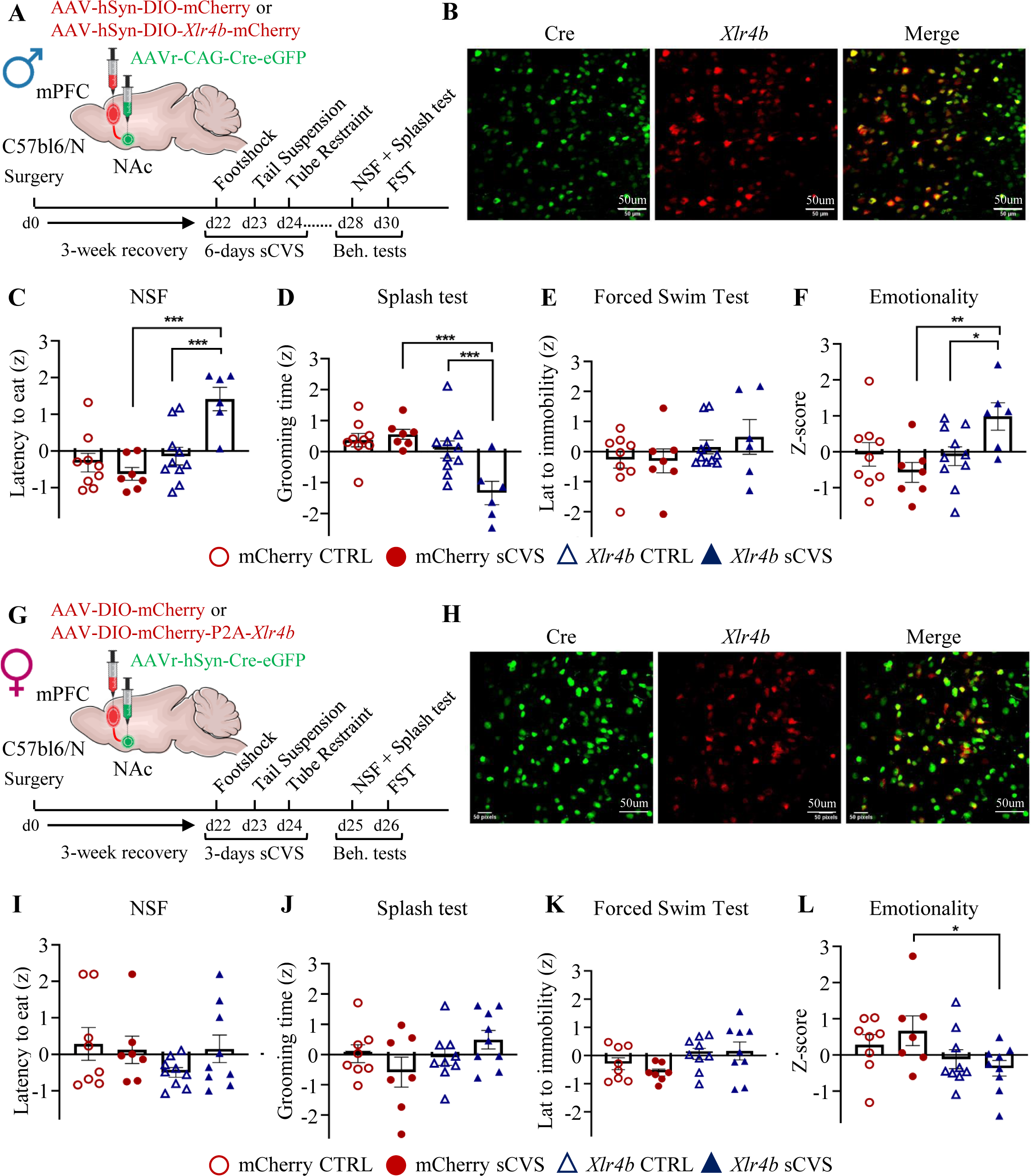
*Xlr4b* overexpression in NAc-projecting mPFC neurons coupled with stress is sufficient to induce anxiety- and despair-like behaviors in male mice. **A**, **G.** Schematic representation of virus injection and experimental design in male and female mice, respectively. **B**, **H.** Representatives of a viral infection in the mPFC with Cre (green) and *Xlr4b* (red). **C-F**, **I-L.** Behavioral tests in males and females after sCVS, respectively, in which *Xlr4b* overexpression in NAc-projecting mPFC neurons increased anxiety-like (**C**) and behavior despair-like behavior in males after stress (**D**), but not in females (**I, J**). *Xlr4b* overexpression coupled with sCVS had no effect on forced swim test. (**E**, **K).** *Xlr4b* overexpression in NAc-projecting mPFC neurons coupled with stress increased overall emotionality in males (**F**) and decreased in females (**L**). Two-way analysis of variance with Tukey multiple comparisons tests was used to determine significance. Bars represent the group average ± SEM. \**p*<0.05, ** *p*<0.01 and \*\*\**p*<0.005. sCVS, subchronic variable stress. NAc, nucleus accumbens. mPFC. medial prefrontal cortex. NSF, novelty-suppressed feeding.

### Functional role of *Xlr4b* on cortico-accumbal mPFC neuron synaptic activity in stressed males

Our results suggest that the overexpression of *Xlr4b* in mPFC neurons from the cortico-accumbal pathway increases stress susceptibility in male but not female mice. Given the role of *Xlr4b* on the development of dendritic spines and the modulation of synapses previously described^72^, we hypothesized that the behavioral effect of *Xlr4b* overexpression could be mediated by altering the functional properties of mPFC neurons projecting to the NAc. To test this, we used a trans-sectional viral approach to overexpress *Xlr4b* in mPFC neurons projecting to the NAc and performed whole-cell patch-clamp recordings (**Figure 4A**). We started by investigating membrane passive properties, such as resting potential, membrane capacitance, and membrane resistance. Our analysis revealed a significant effect of the main factor *viral approach* on resting capacitance (F_(1,95)_=7.434, *p*=0.008), with cells overexpressing *Xlr4b* showing higher capacitance compared to those expressing *mCherry* (**Figure 4B**). However, no significant effect of *stress* nor *viral approach* was observed in resting potential (*p*=0.81 and *p*=0.84, respectively) or membrane resistance (*p*=0.90 and *p*=0.99, respectively; **Figure 4B**). Additionally, we observed a significant effect of stress in the half-width of the first action potential (F_(1,93)_=7.246, *p*=0.008; Tukey *post hoc* test: *mCherry* sCVS vs. *Xlr4b* CTRL, *p*_adj_=0.016). However, there was no significant difference in cell excitability, as indicated by rheobase (*viral approach p*=0.27; *stress p*=0.71). We next subjected mPFC neurons from the cortico-accumbal pathway to steps of increasing injected current. We performed a linear mixed-model analysis with *applied current*, *viral approach*, and *stress* as the fixed effects (**Table 1**). The results show a significant *current* effect (CI95%: [0.06 – 0.07], *p*=<0.001), and a significant interaction between *applied current* and *viral approach* (CI95%: [0.01 – 0.01], *p*=<0.001). We also found a significant difference in mean firing rate between *mCherry* sCVS and *Xlr4b* sCVS, when considering the applied current of 300 pA (Tukey *post hoc* test: *mCherry* sCVS vs. *Xlr4b* CTRL, *p*_adj_=0.009, **Figure 4C**). Overall, this analysis suggests that *Xlr4b* overexpression increases the mean firing frequency when increasing current is applied in NAc-projecting mPFC neurons (**Figure 4C**).

**Figure 4.**
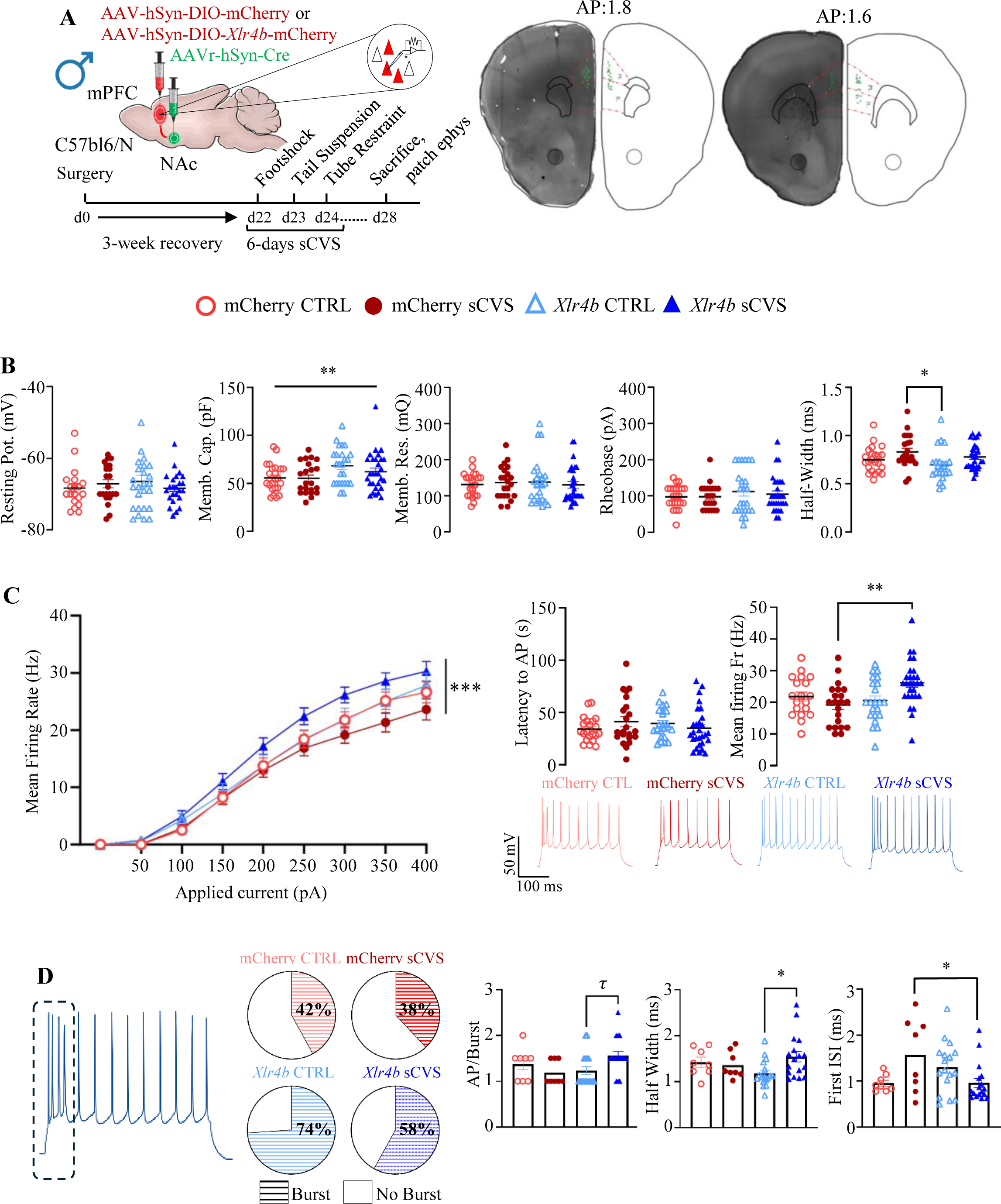
The overexpression of *Xlr4b*, coupled with subchronic stress, increases spiking frequency in response to injected current in cortico-accumbal neurons. **A.** Experimental design. Whole-cell patch-clamp recordings are obtained from *mCherry*-positive neurons and confirmed *post hoc*. Stereotaxic map of recorded cells, as per *post hoc* reconstruction. Each green triangle represents a patched cell. **B.** Passive properties of recorded cells (left, legend code shown above). Excitability and spike properties measurements were obtained from the first observable spike at rheobase (right). **C.** Current-Frequency relation plot. Linear mixed-effects model revealed a significant interaction between *‘applied current’* and *‘viral approach’* (*p*=<0.001). Tukey *post hoc* comparison additionally revealed a significant difference between ‘*mCherry* sCVS’ and ‘*Xlr4b* sCVS’ (*p*_adj_=0.0091). Inset – latency to AP and mean firing frequency at 300 pA (current point picked arbitrarily). Examples of recording traces are provided for each condition. **D.** Burst analysis at 300 pA injected. From left to right: Burst probability: *Xlr4b* OE alone is sufficient to increase the burst probability to 300 pA current injection. Number of APs per burst: the combination of *Xlr4b* manipulation and stress increases the number of action potentials per burst (*p*=0.0523; Tukey *post hoc* comparison); Half-Width: The average burst half-width is increased in the *Xlr4b* sCVS group, compared to *Xlr4b* CTRL (P=0.023; Tukey *post hoc* comparison). No difference is detected between the *mCherry* conditions; Inter-Stimulus-Interval (ISI): The ISI is decreased in the *Xlr4b* sCVS group, compared to *mCherry* sCVS (P=0.042; Tukey *post hoc* comparison). CTRL, control. sCVS, subchronic variable stress.

**Table 1.**
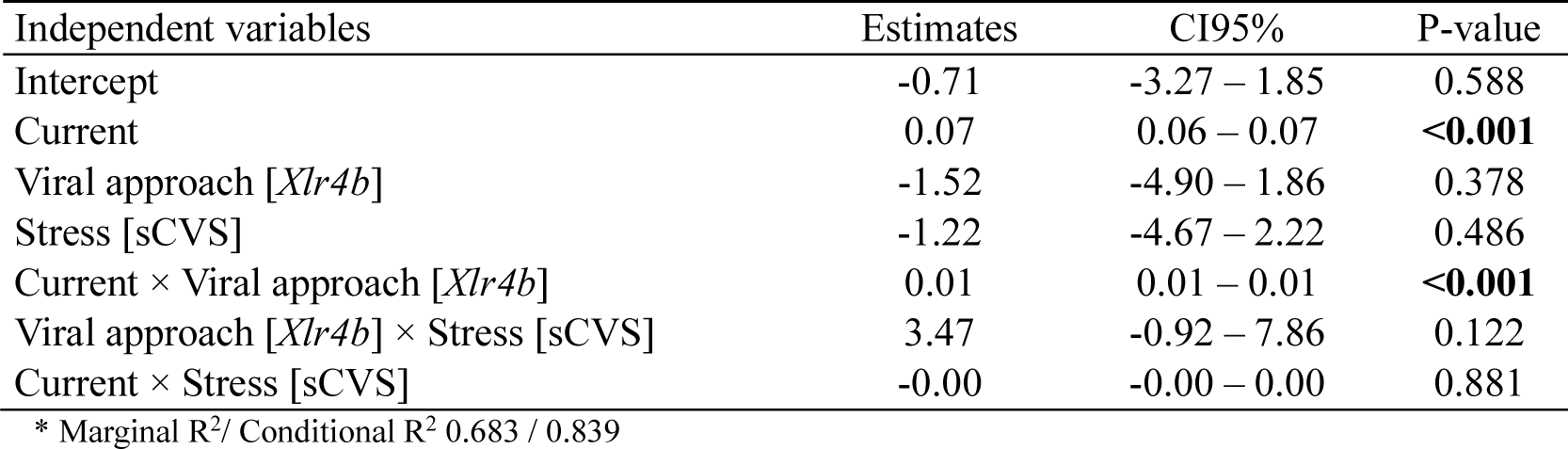
Linear mixed-effects model evaluating the effect of current, viral approach, and stress on mean firing frequency in mPFC neurons.

| Independent variables | Estimates | CI95% | P-value |
| --- | --- | --- | --- |
| Intercept | -0.71 | -3.27 – 1.85 | 0.588 |
| Current | 0.07 | 0.06 – 0.07 | **<0.001** |
| Viral approach [*Xlr4b*] | -1.52 | -4.90 – 1.86 | 0.378 |
| Stress [sCVS] | -1.22 | -4.67 – 2.22 | 0.486 |
| Current × Viral approach [*Xlr4b*] | 0.01 | 0.01 – 0.01 | **<0.001** |
| Viral approach [*Xlr4b*] × Stress [sCVS] | 3.47 | -0.92 – 7.86 | 0.122 |
| Current × Stress [sCVS] | -0.00 | -0.00 – 0.00 | 0.881 |

\* Marginal R<sup>2</sup>/ Conditional R<sup>2</sup> 0.683 / 0.839

Moreover, our results suggest that *Xlr4b* overexpression alone increases the probability of burst firing at high injected currents (**Figure 4D**). The cumulative effect of subchronic stress on *Xlr4b* overexpression shifted the burst activity from a “slow” to a “sustained” burst firing mode. We report a significant *viral approach* × *stress* interaction effect regarding the number of AP/burst (F_(1,46)_=5.591, *p*=0.022), with sCVS *Xlr4b*-expressing cells firing a higher number of action potentials per burst (Tukey *post hoc Xlr4b* CTRL × *Xlr4b* sCVS, *p*_adj_=0.052). We also observed stress-driven differences regarding the duration of the first Inter Spike Interval (ISI) (F_(1,46)_=9.243, *p*=0.003; Tukey *post hoc mCherry* sCVS × *Xlr4b* sCVS, *p*_adj_=0.042). Finally, we considered the half-width of the first AP of each burst to account for kinetic differences induced by stress or transgenes expression. We report a strong trend toward significance for *viral approach* × *stress* interaction (F_(1,46)_=3.903, *p*=0.054) with stress increasing half-width in *Xlr4b* OE cells only (Tukey *post hoc Xlr4b* CTRL × *Xlr4b* sCVS, *p*_adj_=0.02; **Figure 4D**). Together, our results support the hypothesis that *Xlr4b* alters the firing response of NAc-projecting mPFC neurons by changing their passive and functional properties, which ultimately may lead to the development of anxiety-like behavior in male mice.

### Morphological assessment of *Xlr4b* overexpression in stressed male mice

We next hypothesized that *Xlr4b* overexpression alone and paired with sCVS could dysregulate dendritic protein modulation and reduce dendritic arborization as suggested before^72^. We addressed this question using a trans-sectional viral strategy combined with confocal imaging of NAc-projecting pyramidal neurons to perform computational neuronal reconstruction and Sholl analysis (**Figure 5A**). Our analysis shows a significant main effect of *Xlr4b* on overall dendritic arborization (**Figure 5B**, **Table 2**). Specifically, our results suggest that *Xlr4b* overexpression reduces arborization by 24% independently of the distance. Further analysis showed a significant *Gene x Distance* interaction, suggesting that *Xlr4b* overexpression causes a stronger reduction in dendritic complexity in the proximal sections of NAc-projecting mPFC neurons compared to the distal sections (**Figure 5B**). More precisely, with each incremental unit in distance from the soma, the dendritic complexity decreased by an average of 0.86 times (RR=0.86, CI95%: [0.84 – 0.89], *p*<0.001). Overall, *Xlr4b* reduced dendritic complexity by 0.76 times (RR=0.76, CI95%: [0.60 – 0.96], *p*=0.022) compared to the control, accounting for all other random and fixed effects. Interestingly, the significant *Gene x Distance* interaction further indicates that the effect of *Xlr4b* grows stronger with distance, leading to a net increase in complexity at greater distances rather than the expected reduction, with a rate ratio increase of 1.04 per unit distance (RR=1.04, CI95%: [1.01 – 1.08], *p*=0.020). This means that while complexity decreases due to distance and *Xlr4b*, at greater distances, the effect of *Xlr4b* reverses, leading to an increase in complexity rather than a decrease. Finally, we used the same viral approach to evaluate the effect of *Xlr4b* overexpression combined with sCVS on spine density. Our analysis shows the effect of *stress* (F_(1,44)_=13.38, *p*=0.0007) and *Xlr4b* overexpression (F_(1,44)_=10.05, *p*=0.0028) on spine density, exclusively. We found that sCVS alone is sufficient to reduce spine density in male mice (CI95%: [0.0002 – 0.34], *p*_adj_ < 0.049; **Figure 5C**). *Xlr4b* overexpression, by itself, did not change spine density (CI95%: [-0.22 – 0.02], *p*_adj_ < 0.15) although its overexpression in males with sCVS abolished the effect of stress and normalized spine density compared to control mice (CI95%: [-0.10 – 0.14], *p*_adj_ < 0.97). Furthermore, we identified a trend in the decrease of spine density when *Xlr4b* overexpression and sCVS are combined (CI95%: [-0.005 – 0.243], *p*_adj_ < 0.06; **Figure 5C**). Together, this suggests that *Xlr4b* overexpression changes neuronal complexity and spine density.

**Figure 5.**
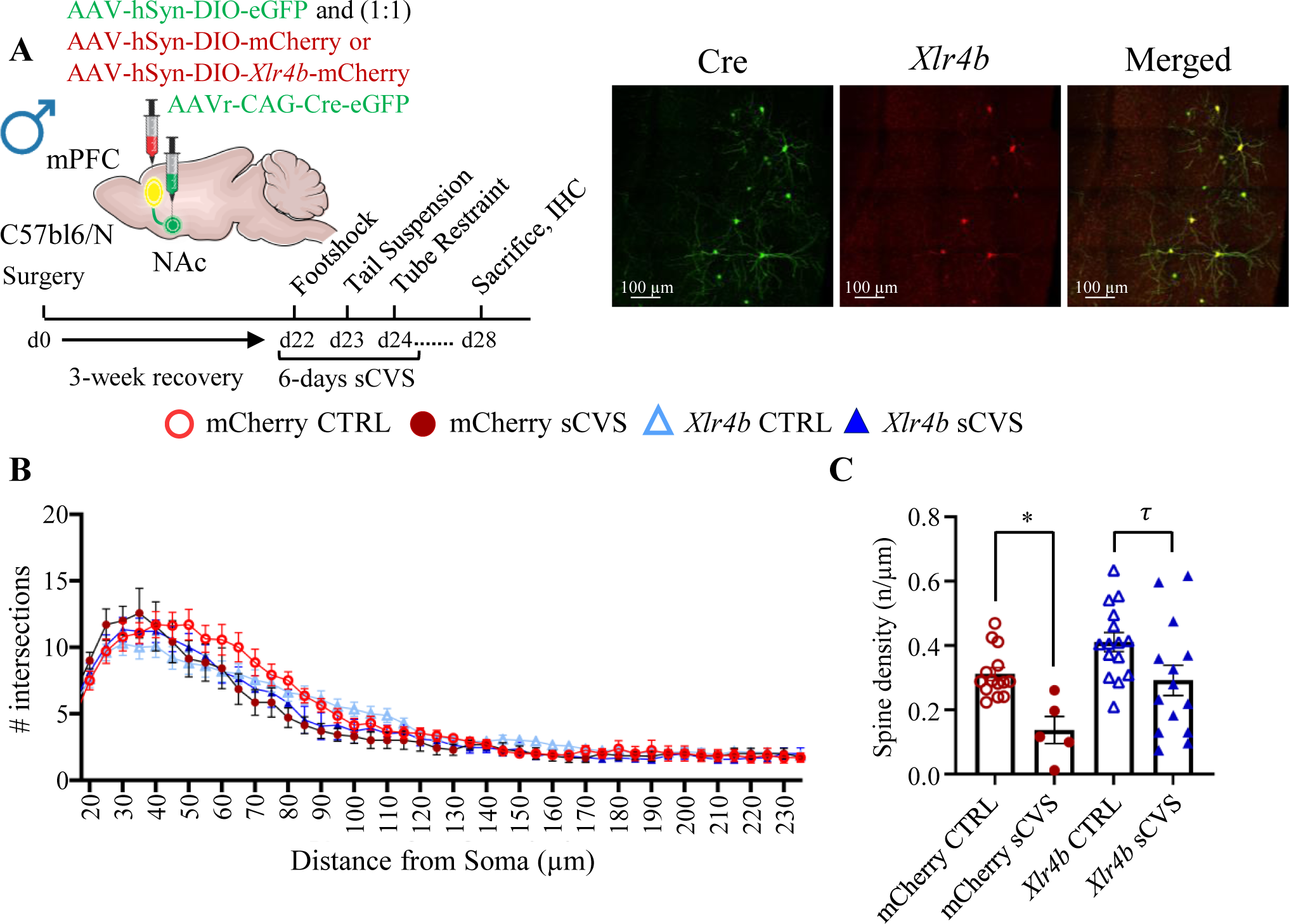
The overexpression of *Xlr4b* and subchronic stress changed morphology and spine density in cortico-accumbal neurons. **A.** Schematic representation of a mouse brain showing the mPFC and NAc pathway connections, the viral injection sites, and the experimental timeline for sCVS, in addition to representative images of a viral infection in the mPFC with *Xlr4b* (green) and *mCherry* (red). **B.** Sholl analysis graph showing the number of intersections and the distance from the soma for the 4 groups, a combination of control, sCVS, and *Xlr4b*. **C.** Two-way analysis of variance with Tukey multiple comparisons tests was used to determine significance. Bars represent the group average ± SEM. τ *p*_adj_<0.1, * *p*_adj_<0.05, and ** *p*_adj_<0.01. CTRL, control. sCVS, subchronic variable stress.

**Table 2.** Random effects (intercept and slopes) model evaluating the effect of distance, condition, and viral approach on NAc-projecting mPFC neuronal dendritic arborization.

| Independent variables | Estimates | CI95% | P-value |
| --- | --- | --- | --- |
| Intercept | 13.34 | 11.24 – 15.83 | **<0.001** |
| Distance | 0.86 | 0.84 – 0.89 | **<0.001** |
| Viral approach [*Xlr4b*] | 0.76 | 0.60 – 0.96 | **0.022** |
| Condition [sCVS] | 0.95 | 0.71 – 1.27 | 0.722 |
| Distance × Viral approach [*Xlr4b*] | 1.04 | 1.01 – 1.08 | **0.020** |
| Distance × Condition [sCVS] | 0.99 | 0.94 – 1.03 | 0.521 |
| Viral approach [*Xlr4b*] × Condition [sCVS] | 1.28 | 0.85 – 1.94 | 0.234 |

\* Marginal R<sup>2</sup>/ Conditional R<sup>2</sup> 0.676/0.881

## Discussion

Stress responses are modulated by the activity of neuronal circuits, the activity of which is under precise transcriptional control. Here, we investigated the impact of stress on the transcriptional organization of mPFC neurons projecting to the NAc as part of the cortico-accumbal pathway. Our analysis identified several gene modules associated with the effect of stress in males and females. We used viral-mediated gene transfer to reproduce the stress-induced expression profile of a key hub gene and confirmed its role in stress susceptibility in a sex-specific fashion. Additionally, we measured the impact of these variations on the activity and morphology of NAc-projecting mPFC neurons revealing important changes underlying the behavioral effects induced by stress. Overall, our findings suggest that stress-induced reorganization of transcriptional structures in NAc-projecting mPFC neurons induces emotional responses in a sex-specific fashion by interfering with the morphological and functional neuronal properties of this neuronal pathway.

Examining the gene expression profiles from NAc-projecting mPFC neurons, we identified specific genes and molecular pathways that are differentially regulated in response to stress, with low overlap with bulk analyses from the mPFC. Network analysis further revealed how these genes interact, forming intricate regulatory networks controlling emotional responses. Our results pointed towards *Xlr4b* as a main driver of stress susceptibility in males. *Xlr4b* is found on the X chromosome and is paternally repressed following a dynamic tissue-specific pattern^77^. It has been shown to have a role in the control of dendrite and spine formation through a Cux1/2-dependent mechanism^72^. Furthermore, it has been associated with working memory and affective deficits in mice^72,73^ and autism and schizophrenia in humans^74–76^. Here, we expand on the role of *Xlr4b* as a driver of stress susceptibility in males. Interestingly, our analysis shows that *Xlr4b* is induced by stress only in males. Indeed, at baseline, this gene is barely expressed in both sexes. However, it acts as a hub gene in its respective module and is significantly associated with stress phenotype in male mice. Interestingly, *Xlr4b* has been suspected to have epigenetic properties through the remodeling of chromatin structures^72^. Once expressed, this gene may impose coordinated control of its network members via a complex reorganization of epigenetic states and, as such, become a major driver of a large proportion of its respective network.

Although the cortico-accumbal pathway has been frequently associated with the expression of stress responses, whether it is potentiated or blunted by chronic stress remains unclear^21–23,78,79^. Here, we provide data suggesting that anxiety and despair-like behaviors are induced by elevated cortico-accumbal activity. It is interesting to note that *Xlr4b* overexpression alone is not enough to elicit such changes, indicating that the behavioral outcome is a consequence of the interaction between gene expression and stressful stimuli. This is consistent with previous work showing that the chemogenetic activation of this pathway induces the expression of anxiety and despair-like behaviors in males and females while its inhibition rescues anxiety and despair-like behaviors in males but not females^26^. It is also in agreement with functional studies showing that continuous stimulation of the mPFC nerve terminals in the NAc decreases social preference^21^ and hyperexcitability of the NAc-projecting mPFC neurons reduces reward-motivated behaviors in male rats^22^. Interestingly, our transcriptional screening identified key players underlying these effects including *Xlr4b*. The overexpression of this gene resulted in a significant reconfiguration of cortico-accumbal neurons’ firing properties. Indeed, cells overexpressing *Xlr4b* exhibited a larger firing rate upon increasing applied currents implying pronounced cellular excitability of NAc-projecting mPFC neurons. Interestingly, we previously showed that CVS increases excitatory inputs on mPFC neurons projecting to the NAc. Thus, the sum of these converging excitatory inputs is likely to increase the current flow going through NAc-projecting mPFC neurons and, combined with the stress-induced overexpression of *Xlr4b*, results in general hyperactivity of this pathway in stressed mice.

Another way *Xlr4b* overexpression may impact neuronal activity and change stress susceptibility is by imposing a morphological reconstruction of NAc-projecting mPFC neurons. Indeed, morphological changes in the mPFC have consistently been reported in humans with MDD and mouse models of chronic stress^8–10^. We have also shown that stress significantly reduces the arborization of NAc-projecting mPFC neurons^26^. Here, our results suggest that, at a short distance from the soma, overexpression of *Xlr4b* decreased neuronal complexity. Our results also show a trending decrease in spine density induced by *Xlr4b* overexpression following subchronic stress, which is consistent with previous reports on *Xlr4b* functions^72^. Fewer spines could contribute to an increase in neuronal excitability by allowing proximal inputs to have a more direct and potent impact on triggering the action potential^80^. This could also lead to compensatory changes in neurons, such as increased synaptic strength via modulation of receptor density, making them more responsive to stimuli^81^. Furthermore, spine position is also critical, as spines located closer to the soma are less subjected to attenuation. Globally, this suggests that *Xlr4b* overexpression in stressed mice could promote heightened excitability, resulting in a higher firing rate despite reduced structural integration.

Besides, our findings pointed toward a substantial increase in burst-firing activity in *Xlr4b*-overexpressing neurons. Interestingly, the combination of sCVS with *Xlr4b* overexpression resulted in a transition from slow to high burst activity. This transition is supported by the elevated number of action potentials per burst with an increased half-width of the burst spikes, as described before^82^. The oversimplified dendritic arborization seen in *Xlr4b*-overexpressing cells could lead to a reduction of the firing threshold, enhancing burst patterns, which would result from the elevated frequency of excitatory inputs on NAc-projecting mPFC neurons observed before^26^. In turn, elevated activity would change the density of calcium voltage-gated channels and receptors, which would further amplify neuronal excitability, promoting high-frequency firing patterns and further potentiating neuronal excitability^80,81^.

It is important to mention the limitations intrinsically associated with the RiboTag approach used in this study^44^. Indeed, the gene content analyzed in this study may not be specific to the NAc-projecting mPFC neurons and should rather be considered enriched in this neuronal pathway. To compensate for this aspect, we employed a series of filters to remove non-neuronal genes based on previously published mouse cell type-specific datasets^50^. Although this does not ensure cellular specificity, it does mitigate these limitations by providing transcriptional profiles enriched in this neuronal pathway. Another important point refers to the challenges posed by biological sex when comparing male and female transcriptional profiles. Indeed, several biological factors such as genes expressed on the sexual chromosomes, sexual hormones, epigenetic mechanisms, and a series of transcription factors can modulate transcriptional profiles and the molecular mechanisms underlying stress responses in males and females differently^83–86^. While we adopted a series of statistical approaches to compensate for these effects at the transcriptional level, future work should implement experimental designs aiming at addressing these questions more in detail to eventually delineate a more refined landscape of sex differences.

To conclude, chronic stress changes the activity of the mPFC and more precisely, its circuitry, including the cortico-accumbal pathway. These functional changes result in the expression of maladapted emotional responses. Here, we mapped the transcriptional signatures underlying these functional and behavioral changes in a sex-specific fashion. We identified gene signatures associated with a reorganization of the functional and morphological properties of NAc-projecting mPFC neurons. Globally, this provides further evidence of the molecular mechanisms underlying the expression of stress responses in males and females while further strengthening the contribution of the cortico-accumbal pathway in emotional responses to chronic stress. Further work should investigate these effects in other neuronal pathways within the mPFC and evaluate the consequences of the molecular and functional changes described in this study on synaptic communication in downstream structures. Nevertheless, by describing the gene signatures associated with chronic stress in the cortico-accumbal pathway, our findings provide a molecular substrate on which to act to modulate the activity of this neuronal pathway and potentially counteract some of the detrimental consequences of chronic stress at the morphological, functional and behavioral levels in males and females.

## Acknowledgments

We would like to thank the *Plateforme d’Outils Moléculaires* (https://neurophotonics.ca/fr/pom) at CERVO Brain Research Centre for viral vector production. We also thank Thibault Bittar for his help in the early phase of the project. BL holds a Sentinelle Nord Research Chair, is supported by the Canadian Institutes of Health Research (Grant No. PJT-451728 and PJT-451858), and the Natural Science and Engineering Research Council of Canada (Grant No. RGPIN-2019-06496) and receives Fonds de Recherche en Santé du Québec (FRQS) Junior-2 salary support. AMP, LP, KH and AMR are supported by FRQS scholarships. CDP lab is supported by CIHR grant PJT169117 and NSERC grant RGPIN-2017-06131. ML Lab is supported by a CIHR grant 451548 and NSERC grant RGPIN-2024-05363. ML also receives FRQS-Parkinson Quebec Senior salary support.

## Author contributions

B.L. conceived the project, designed the experiments, and wrote the manuscript. ML and CDP contributed to the experimental design and reviewed the manuscript. ML also provided the RiboTag animals and contributed to the RiboTag experimental design. AMP, SM and AMR performed the bioinformatic analyses. LBA performed the behavioral analyses. LP performed the electrophysiological experiments. MdA and KH performed the morphological analysis. All authors contributed to the preparation of the manuscript.

---

**Supplemental Figure 1.**
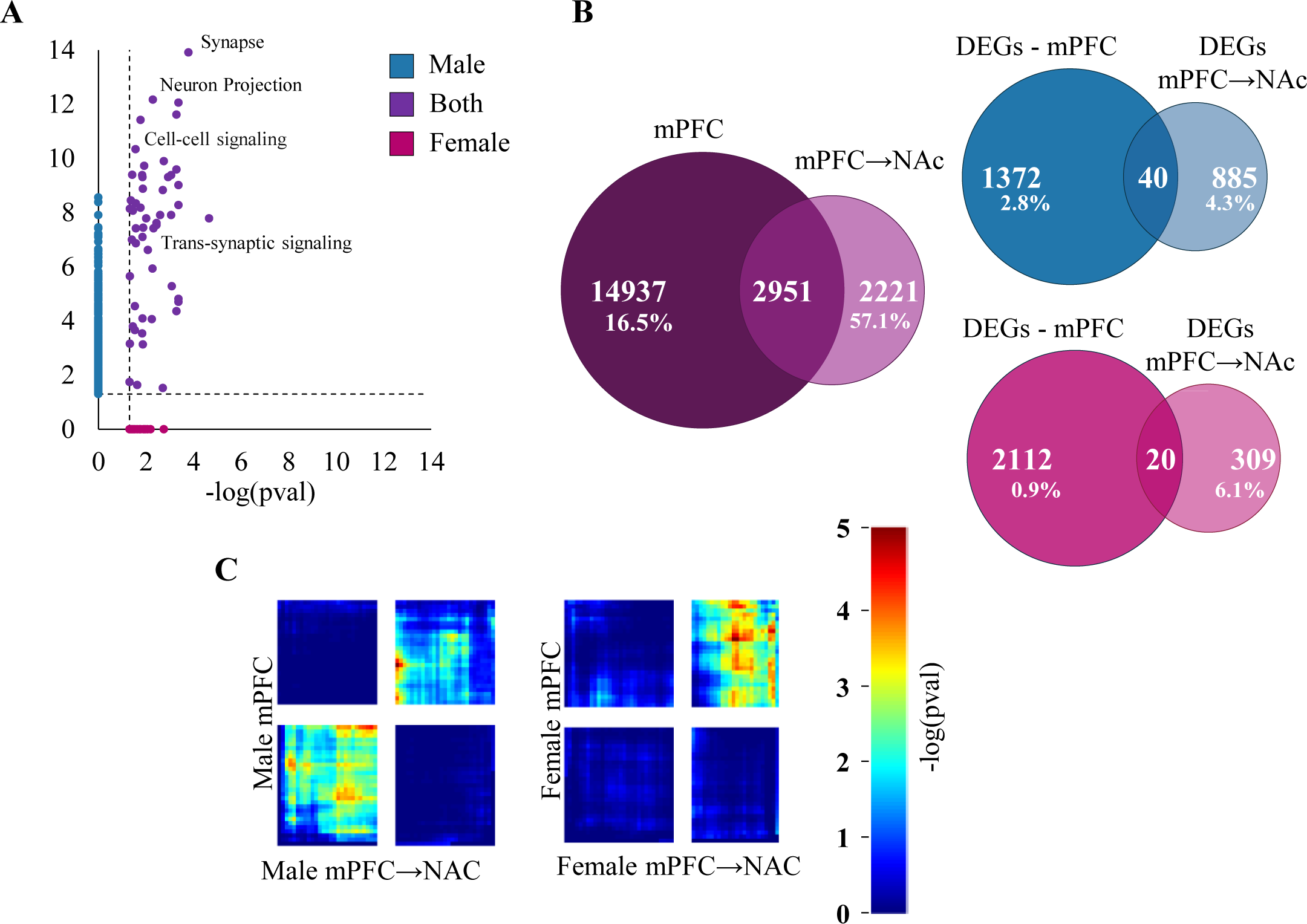
**A.** Scatter plot showing common (purple) or exclusive (blue for males and pink for females) gene ontology terms for biological process, cellular component, and molecular function, combined. The dotted line represents *p*=0.05. **B.** Venn diagrams showing the intersection between bulk RNA sequencing of the whole mPFC and the mPFC-to-NAc pathway, for genes included in the analysis for mPFC (purple) and the mPFC-to-NAc pathway (light purple), and DEGs (*p* ≤ 0.05) for males and females (blue and pink, respectively). They indicate low transcriptional overlap between all comparisons for mice after CVS, while highlighting unique genes on the mPFC-to-NAc pathway that were not expressed in the bulk mPFC analysis. **C.** RRHO heatmaps showing threshold-free transcriptional overlap between bulk RNA sequencing of the whole mPFC and the mPFC-to-NAc pathway for male (up) and female (down) mice after CVS. The lower left and upper right quadrants show a low-intensity overlap for similarly upregulated and downregulated genes, respectively, except for females, with almost no overlap on the commonly upregulated genes.

**Supplemental Figure 2.**
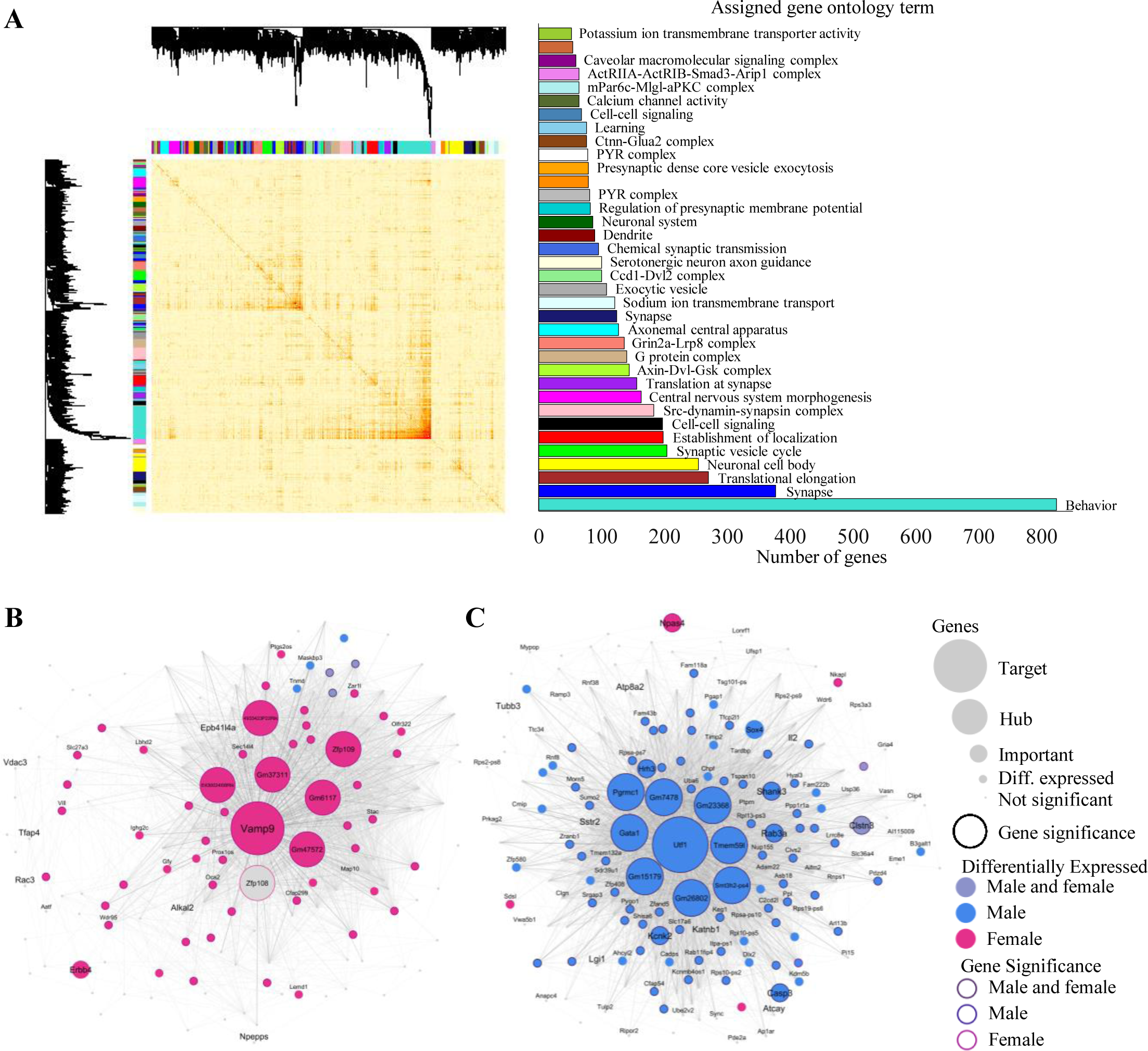
**A.** Topological overlap matrix plot for all samples included in the analysis, in which light color represents low topological overlap, and darker color represents high overlap. The bar plot represents the number of genes alocated to each gene module, whith their top ranking assinged gene ontology term, if available. **B, C.** Cytoscape representation of the *Magenta* (most relevant module in females, not relevant in males) and *Black* (third most relevant module in males, not relevant in females) modules, respectively, representing genes differently based on their connectivity, expression and significance.

