## Supplemental Table 1 for "Transcriptional profiling of the cortico-accumbal pathway reveals sex-specific alterations underlying stress susceptibility"

**Supplemental Table 1.** Adeno-associated viral vectors

| **Serotype** | **Promoter** | **Construct** | **Provider** | **Lot** | **GC/mL** |
|---|---|---|---|---|---|
| AAV2 retro | hSyn | cre | POM | 845 | 2.10E+13 |
| AAV2 retro | CAG | cre-eGFP | POM | 2350 | 1.10E+13 |
| AAV2/5 | hSyn | DIO-*Xlr4b*-GSG-T2A-*mCherry* | POM | 1831 | 1.60E+13 |
| AAV5 | hSyn | DIO-*mCherry* | POM | 2350 | 1.10E+13 |
| AAV5 | hSyn | DIO-eGFP | Addgene | V54234 | 1.30E+13 |

Abbreviations: AAV, adeno-associated virus; DIO, double-inverted orientation; eGFP, enhanced green fluorescent protein; POM, CERVO Brain Research Centre's *Plateforme d'Outils Moléculaires*.
